# RACLET: the Ramp Above Critical Level Endurance Test to evaluate critical velocity in running

**DOI:** 10.1101/2025.07.11.664350

**Authors:** Mylène Vonderscher, Pierre Samozino, Maximilien Bowen, Baptiste Morel

## Abstract

Critical velocity (*V*_*c*_) is an important fatigability threshold in running, but only strenuous methods exist to assess it (e.g. time-trials). Based on a novel mathematical model of fatigability, the Ramp Above Critical Level Endurance Test (RACLET), a simple, non-exhaustive, 5-min test to evaluate *V*_*c*_ was developed. This study aimed to test the reliability and validity of the RACLET. Thirty-eight participants performed two RACLET (session 1) and three time-trials (sessions 2-4). For the RACLET, velocity target tracking was guaranteed by either a pacing bike or cones and whistle signals. GPS was used to measure the participants’ running velocities. *V*_*c*_, *τ*, and 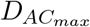 were determined by adjusting the fatigability model on the measured *V*_*max*_. The RACLET test-retest and RACLET vs. time-trials parameters were compared for reliability and validity testing, respectively. The RACLET *V*_*c*_ is reliable and valid (mean difference=-1.7*±*1.9% and 6.2*±*6.5%, respectively). Although *τ* and 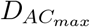 presented important variability (s.d. difference ~30%), the combination of the RACLET parameters enabled an excellent prediction of time-trials performance (mean error=-8.9*±*5.1%, 0.9*±*7.4%, −2.0*±*5.2% for 400, 1500, and 3000 m, respectively). RACLET is a novel non-exhausting test that enables a reliable and valid assessment of the velocity–time relationship parameters.

## Introduction

Running performance is highly dependent on the ability to sustain high intensity over time. The relationship between a given intensity and the time it can be maintained was first described in animals by Kennelly (1906). It was then applied to human performance by Hill (1925) before being mathematically formalised with an inverse function by Monod and Scherrer (1965). The intensity-time relationship is well known through the Critical Power (P_*c*_, in W) concept, but will be described hereafter in terms of velocity as preferred when focusing on linear running. The velocity–time (*V* (*t*)) relationship corresponds mathematically to a hyperbolic function defined by two parameters: the asymptote, namely, the critical velocity (*V*_*c*_, in m ·s^*−*1^) and the curvature constant or “distance capacity” (D*′*, in m). The last two parameters are analogous to P_*c*_ and the mechanical energy reserve (W*′*, in J) respectively (Housh et al., 1991; Pepper et al., 1992). *V*_*c*_ is defined as the boundary between the heavy and severe intensity domains (Burnley et al., 2012). This implies that running at a slower velocity than *V*_*c*_ enables a physiologically stable state (in *V O*_2_, Gaesser and Poole, 1996; in blood flow, Copp et al., 2010; or in intramuscular concentration of inorganic phosphate, Jones et al., 2008). Conversely, exercising above *V*_*c*_ (i.e. in the severe intensity domain) generates drastic fatigability (defined as the loss in maximal capacities resulting from muscle activity and reversible by rest, Gandevia, 2001). In the severe intensity domain, the finite distance reserve D*′* available at the beginning of an exercise is depleted. The total depletion of D*′* results in exhaustion (i.e. the end of the task or a decrease in intensity to *V*_*c*_). The concept of *V*_*c*_ is of great interest and has been used in a wide range of fields (e.g. in patients for rehabilitation (Dolmage et al., 2012) or athletes for training (Meyler et al., 2025) and pacing purposes (Pettitt, 2016). However, the currently available methods that aim to assess *V*_*c*_ are extremely strenuous.

Traditionally, *V*_*c*_ has been obtained from multiple tests of different intensities, leading to exhaustion. Historical methods include either the completion of multiple trials at various fixed velocities to sustain for as long as possible until exhaustion (time to exhaustion, TTE, Hughson et al., 1984; Pepper et al., 1992) or various fixed distances to cover within the shortest duration (time-trials, TT, Galbraith et al., 2014; Kordi et al., 2019). More recently, *V*_*c*_ was derived from a single 3-min all-out test (Black et al., 2014; Pettitt et al., 2012). Although less time-consuming, this method is still extremely strenuous. All of the above-mentioned methods require participants to be highly motivated. Indeed, it is well known that perception of effort influences performance during such tests (Salam et al., 2018). Therefore, determining *V*_*c*_ remains a challenge. Other promising methods using raw *in situ* training data (Smyth and Muniz-Pumares, 2020) that mimic the power record profile concept in cycling (Pinot and Grappe, 2011) have been developed. Although probably more representative of *in natura* conditions, the exercise is less controlled than in a laboratory environment. In addition, a large amount of data must be collected to establish an athlete’s profile, making it difficult to monitor progress. Finally, the self-determination of *V*_*c*_ along a 10-min submaximal test has recently been validated (Follador et al., 2020). Although this helps overcome the motivational limitation, this method does not provide any information about D*′*. It is also probable that the perceived effort is no longer linked to *V*_*c*_ when the experimental conditions are modified (e.g. in a hot environment).

A recently proposed integrative model can overcome these methodological limitations. Based on the pioneering work of Morton (1994), Bowen et al. (2024) proposed a generic formulation for exercise fatigability in the severe domain under isokinetic conditions. This model explains fatigability through a proportional relationship between the alteration of maximal capacities and the exercise intensity integral performed above the critical intensity. Commonly expressed in terms of power, the concept of critical intensity can be adapted in running to describe the alteration of maximal velocity capacities over any fatiguing exercise in the severe domain (when V>*V*_*c*_) under iso-resistance conditions (e.g. unconstrained level running):

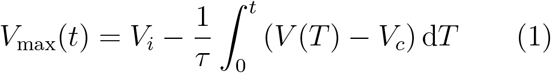

where *V*_*i*_ is the initial maximum velocity (in m · s^*−*1^) that can be reached before the onset of any fatigability, 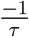 is the proportionality coefficient (in s^*−*1^) between the distance accumulated above *V*_*c*_ (also hereafter referred to as *D*_AC_) and the decrease in maximal capacities, *V* is the velocity target corresponding to the fatiguing task (which can change over time; in m · s^*−*1^) and *V*_max_ is the maximal velocity (in m · s^*−*1^). Note that D*′* is further referred to as 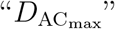 with 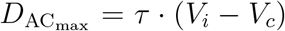. According to the original formalism, *D*_AC_ corresponds to an absolute portion of 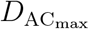 performed over time.

This model has already demonstrated an excellent description of exercise fatigability in the severe domain (Bowen et al., 2024), and has been more recently supported by experimental data from the *adductor pollicis* muscle (Vonderscher et al., 2024). Interestingly, this model has dual uses. First, if the three individual parameters *V*_*i*_, *V*_*c*_, and *τ* as well as the exercise velocity *V* (*t*) are known (with *V* (*t*) *> V*_*c*_), it is possible to predict the magnitude of fatigue induced over time (*V*_max_(*t*)). Second, adjusting the model to observe the maximal capacities in various states of fatigability should enable the determination of the three physiological parameters *V*_*i*_, *V*_*c*_, and *τ*. The novelty of this model lies in its applicability to any exercise performed in the severe intensity domain; the alteration of maximal capacities (fatigability kinetics) being proportionally linked with the accumulation of *D*_AC_, itself directly depending on the exercise target intensity.

Based on this property, a decreasing ramp function appears to be a remarkable target for determining an individual’s *V*_*c*_. Indeed, it can be designed to cover both severe- and heavy-intensity domains, causing a quick onset of fatigability at the beginning of the exercise (i.e. a decrease in maximal velocity capacities; Burnley et al., 2012) and enabling a recovery phase at the end of the test (Skiba et al., 2012, 2015). Conceptually, the precise intensity when a shift between fatigability and recovery occurs corresponds to *V*_*c*_. The Ramp Above Critical Level Endurance Test (RACLET) was developed based on this simple idea and the previously proposed mathematical framework.

No simple method has hitherto been proposed to assess the velocity-time relationship and its related parameters in running, within a unique test that does not lead to exhaustion or maximal fatigability. This study aimed to test the RACLET feasibility, intra-session reliability, concurrent agreement with the *gold standard* method, and running performance prediction capacities. A “hard” but not maximal level of perceived exertion was expected in the most fatigued part of the RACLET, while the active recovery of the last few minutes was expected to deliver a moderate level of perceived exertion immediately after the RACLET. Given the encouraging results already obtained with the novel fatigability model and the rationale for using a decreasing ramp to assess the individual parameters *V*_*i*_, *V*_*c*_, and *τ*, the RACLET was expected to be reliable and valid when compared to the historical time-trials method.

## Methods

### Theoretical background

The RACLET consists of a 5-min decreasing-intensity ramp starting at a velocity *V* ^⋆^ (*V*_*c*_ *< V* ^⋆^ ≤ *V*_*i*_, i.e. in the severe intensity domain) and progressively decreases with a slope (S, in m · s^*−*2^). The end-test velocity (*V*_end_ in m · s^*−*1^) is set *a priori* such that *< V*_*c*_ (i.e. in the heavy or moderate domain). Typically, in an active population, *V*_end_ corresponds to the slowest velocity at which it is still more natural to run than to walk (i.e. 2.1 m · s^*−*1^, Segers et al., 2006). Regularly during the test, *V*_max_ is assessed by short all-out 3-s sprints. From Eq. 2, if *v*(*t*) follows a first-order polynomial function, the fatigability kinetics follows a second-order polynomial function:

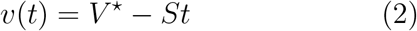

Indeed, based on Bowen et al., 2024, the integrative force–time model and the relationship between force and velocity *F* (*V*):

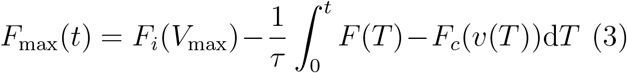

In the case of isotonic running conditions (*F* = cst), the initial (*i*) and critical (*c*) Force-Velocity *F* (*V*) relationships are defined by *F*_0_ (the maximal force at a null velocity) and *V*_0_ (the maximal velocity up to which it is possible to produce force), such as

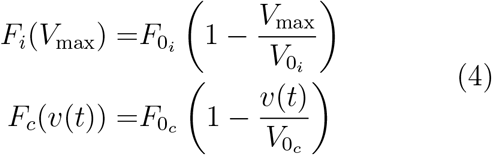

Combining the Eqs. 2, 3, and 4 provides Eq. 5 from which it is possible to determine the evolution of maximal velocity over time (Eq. 6):

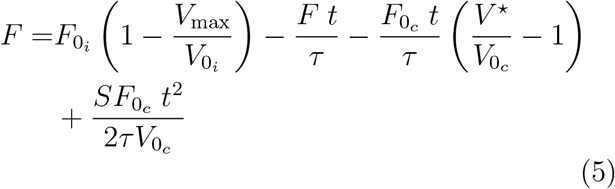

In the particular isotonic running condition (i.e. level ground, all other friction forces being neglected, *F* = 0), by a trivial development, the maximal velocity is defined by:

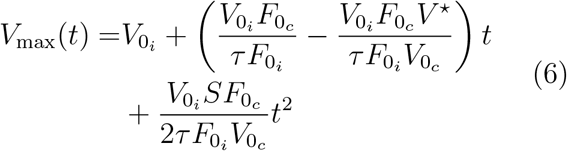

From Eq. 6 and by introducing *α* as the ratio of equivalence between force and velocity parameters 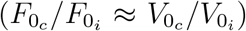, the maximal velocity *V*_max_(*t*) can be defined by a simple second order polynomial function:

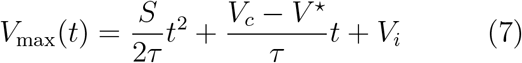

Or more simply:

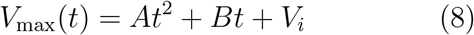

With:

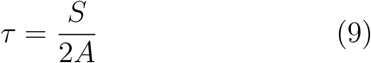

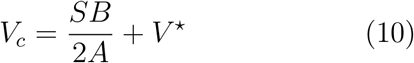

It should be noted that 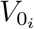 and 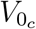 are simplified as *V*_*i*_ and *V*_*c*_, respectively.

### Participants and experimental design

Thirty-eight participants (26 men and 12 women; 21.4 ± 1.3 years old; 66.5 ± 9.2 kg; 173.6 ± 8.3 cm), free of any recent injury, participated in this study. This study was conducted in accordance with the principles outlined in the Declaration of Helsinki for research involving human participants. All participants provided informed consent prior to inclusion in the study. Approval for this project was obtained from the local Ethics Committee on Human Experimentation.

Four sessions were conducted on a 400-m tartan athletic track, at least 24 hours apart. All sessions started with a standardised 30-min warm-up, including 10 min of moderate-intensity running, athletic drills, and progressive sprints.

#### Session 1: RACLET

The first session consisted of two 40-m sprints followed by a submaximal 800-m ran at a self-selected intensity (details are provided below). Participants then performed two identical RACLET (RACLET (A) and RACLET (B)) separated by a 20-min rest period for reliability testing. As mentioned previously, setting up a RACLET requires two specific attention points. First, the participant had to run at a pace following a linearly decreasing ramp, inducing fatigability at the beginning and enabling recovery in the second half of the test. To do so, two protocols were used. The first setup (bike) consisted of following a pacing bike equipped with a speedometer (hall effect sensor and wheel magnet) to respect the target speed. In the second setup (cones), cones were placed every 20 m on the track, and participants self-adjusted their velocity to cover 20 m between two whistle signals. The time intervals between the two whistle signals were adjusted based on the desired target velocity. In both scenarios, the ending velocity was fixed at 2.1 m · s^*−*1^ and the test duration was fixed at 5 min for all participants. Conversely, the starting velocity (*V* ^⋆^) was individualised to induce sufficient fatigability (*V*_max_ alteration) without reaching exhaustion (complete 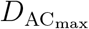 depletion). To set *V* ^⋆^: *V*_*i*_, *V*_*c*_, and *τ* were roughly *a priori* estimated. The mean velocity reached in the last 10-m of the two 40-m sprints was considered as a rough approximation of *V*_*i*_. Participants then ran an 800-m trial during which they were asked to run at the “hardest pace possible to maintain around 40 minutes”. The mean velocity required to complete the second lap (400 m) was considered a rough estimation of *V*_*c*_. Finally, the *a priori* rough estimation of *τ* was fixed at 40 s based on typical data (Kramer et al., 2020). Provided these *a priori* parameter estimations, and using Eq. 7, *V* ^⋆^ was set such that the maximal *D*_AC_ accumulated during the test would theoretically reach 50% of the 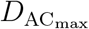. The time at which V was equivalent to *V*_*c*_ was between 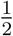 and 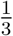 of the test duration.

Second, the maximal velocity capacity of the participants had to be evaluated regularly to provide observations of its decrease (or increase) and to enable the model to be adjusted (and thus determine *V*_*c*_ and *τ*). To ensure a sufficiently thorough coverage of the fatigability phenomenon, participants were asked to perform 3-s all-out sprints at 10 s after the beginning of the test and then every 30 s during the 5-min test (Fig. 1). In the bike setup case, the pacing bike shifted on the side to enable participants to accelerate during the sprint phases. Immediately thereafter, participants were asked to decelerate to follow the pacing bike again. For the cones setup, the participants accelerated for 3 s to reach V_max_ and then slowed down to the target velocity. The sprint phase was not necessarily matching a 20-m distance (e.g. if 35 m had been covered during the sprint, the whistle signal had been blown 5 m before the next cone). In this particular case, participants were asked to maintain the same gap with each following cone for the next 30 s (i.e. a 5-m gap with each cone until the next sprint).

**Figure 1.**
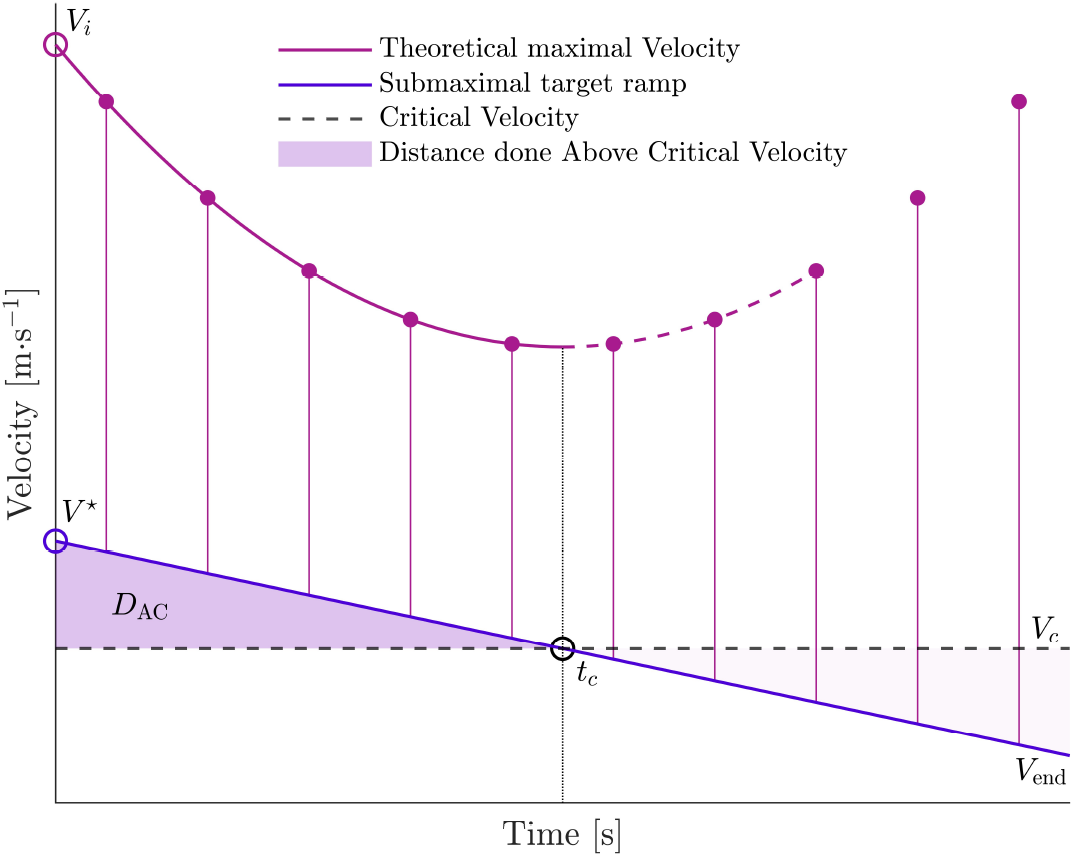
Schematic representation of the Ramp Above Critical Level Endurance Test (RACLET). *V*_*i*_: Initial Velocity, *V*_*c*_: Critical Velocity, *D*_AC_: Distance Above Critical velocity (portion of 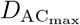): Ramp starting velocity, V_end_: Ramp ending velocity (2.1 *m*·*s*^*−*1^), t_*c*_: Time at which the ramp velocity cross *V*_*c*_ (switch time between fatigability, *i*.*e*. decrease in maximal capacities, and recovery, *i*.*e*. increase in maximal capacities).

Prior to RACLET (A), the participants were familiarised with the procedures by performing the first minute of a RACLET starting at their own *V* ^⋆^ minus 0.5 m · s^*−*1^. A second familiarisation was performed at the found *V* ^⋆^ and stopped after 43 seconds (end of the second sprints). During the two familiarisation, the athletes were asked to “accelerate but not all-out sprint” to limit fatigability prior to the tests. A rest period of at least 5-min was taken before starting the RACLET (A). Participants were equipped with a GPS (GPexe, GK, Udine, Italy) measuring velocity (18 Hz) during the entire session duration (Fig. 2).

**Figure 2.**
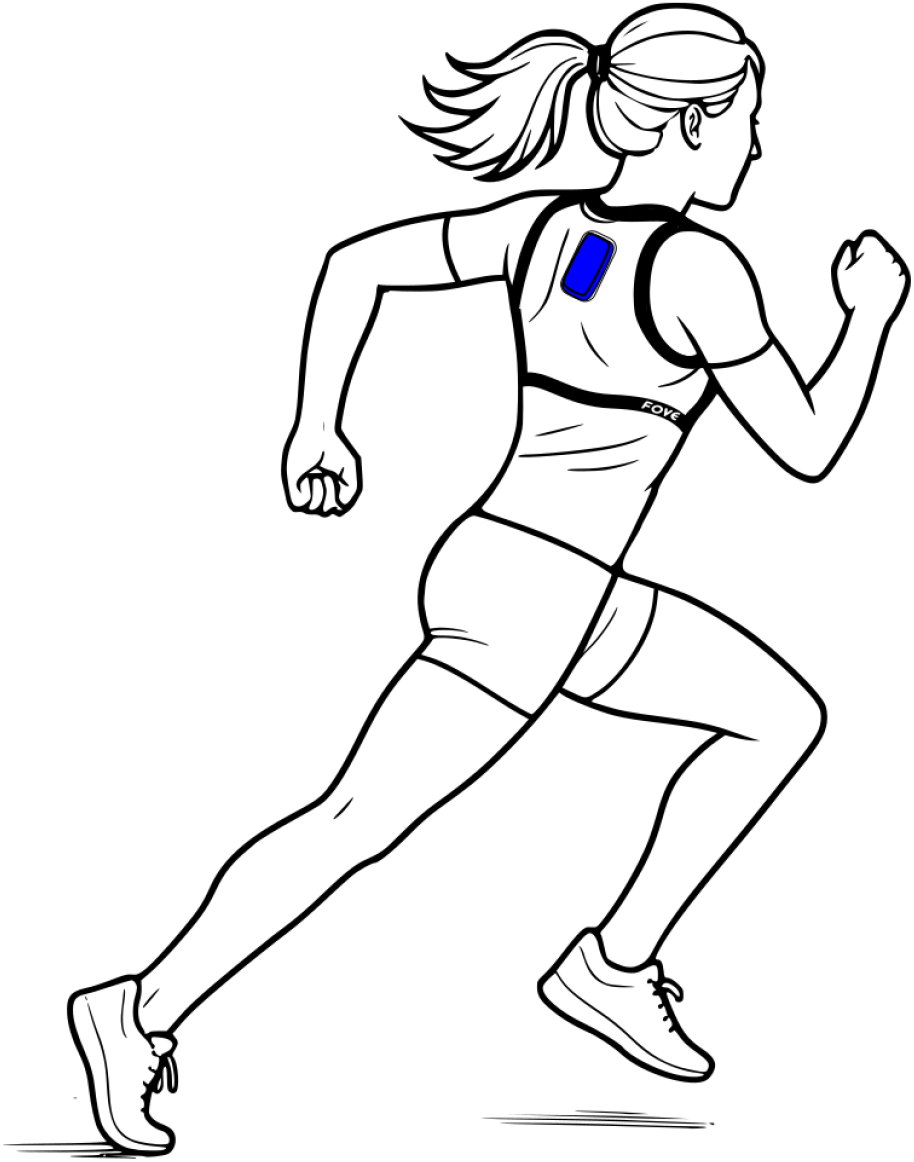
Schematic representation of a participant equipped with the GPS device. In blue: GPexe device in the back of the runner.

To test feasibility, participants were familiarised with the 6-20 Borg scale prior to the RACLET at the beginning of session 1. After each RACLET, participants were asked to rate their perceived exertion “at the most difficult time during the test” and “immediately at the end of the test”.

#### Session 2 to 4: time trials

Sessions 2 to 4 consisted of three time-trials (TT) on 400, 1500, and 3000 m (one TT by session) completed in a randomised order to determine individual velocity-time relationship and then test the RACLET concurrent agreement. Participants were asked to cover each distance within the shortest possible time and adopt the pacing strategy of their choice. The mean running velocity of each trial was then computed using the time required to cover each distance.

## Data analysis

The data were processed using MATLAB (2024b, MathWorks, Natick, USA). The GPS raw velocity data were filtered using a Butterworth low-pass filter (cutoff frequency set at 1 Hz, order 3). The maximum velocity of the best of the two sprints performed after the warm-up was saved as *V*_*i*_. Maximal velocity (*V*_max_) reached during each sprint of RACLET (A) and RACLET (B) were detected. The actual *V* ^⋆^ and *S* of the exercise ramp performed were determined by adjusting Eq. 2 (least square method) on the mean running velocity between each sprint *V*_max_. For each RACLET, all the sprints until the sprint associated with the slowest V_max_ value and the three following sprints were considered in the analysis. The values of V_max_ were filtered using a Butterworth low-pass filter (cutoff frequency set at 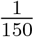 Hz, Order 3). *V*_*c*_ and *τ* were obtained by adjusting Eq. 8 on filtered *V*_max_ values using the least square method (with *V*_*i*_, *V* ^⋆^ and *S* fixed at previously found values; Eqs. 9 and 10).

For the reference method, a homographic function (Eq. 11, Bowen et al., 2024) was adjusted with the least square method on the 3 TT mean velocity and the maximal velocity (*V*_*i*_) detected on the initial sprint that was added as a fourth point at t = 3 s.

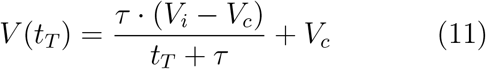

With *t*_*T*_, the time required to cover each distance. For both the RACLET and TT methods, 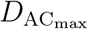 was computed as 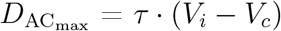.

## Statistical analysis

Absolute (systematic and random differences) and relative reliability were tested on RACLET parameters 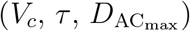 through the change in the mean between RACLET (A) and RACLET (B), the Typical Error of Measurement (TEM), and the Intraclass Correlation Coefficient (ICC), respectively. The concurrent agreement between the two methods, systematic (mean) and random (s.d.) errors between RACLET (A) and TT, were computed for each parameter. RACLET (A) and RACLET (B), as long as the RACLET (A) and TT parameters were compared using a paired-sample t-test after checking for normality. The times to perform 400, 1500, and 3000 m were also predicted using the model of Bowen et al., 2024, from the RACLET (A) parameters (Eq. 12, obtained from Eq. 11) and compared to performances measured during TT through systematic (mean) and random (s.d.) errors, correlation analysis and a paired-sample t-test after checking for normality. All reliability and validity analyses were performed on the entire dataset (pooled bike and cone setups) and on the bike and cones setups separately. The differences in reliability (change in the mean) and validity (systematic error) between bike and cones were tested with an independent-sample t-test for all parameters (*V*_*c*_, *τ*, and 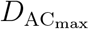). An independent-sample t-test was also conducted to compare the RPE results between the bike and cones modalities. Cohen’s d was used to quantify the effect size and interpreted as small, moderate, large, very large and extremely large effect for values of 0.2, 0.6, 1.2, 2.0 and 4.0, according to Hopkins et al. (2009). The results are expressed as the group mean ± s.d. The significance level was set at p<0.05.

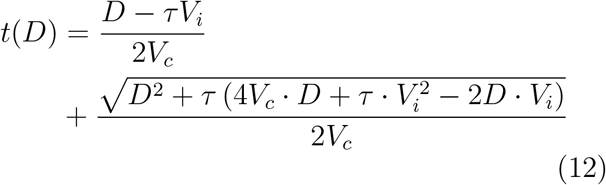

## Results

Among the 38 recruited participants, 19 participants completed session 1 only (with bike setup) and 19 others completed the four sessions (bike, n = 10; cones, n = 9). The RACLET data of 6 tests led to outlier parameter values (either *V*_*c*_ or *τ*), involving 32 participants in the reliability analysis and 16 participants in the validity testing. Typical traces of RACLET (A), RACLET (B), and TT are depicted in Fig. 3.

**Figure 3.**
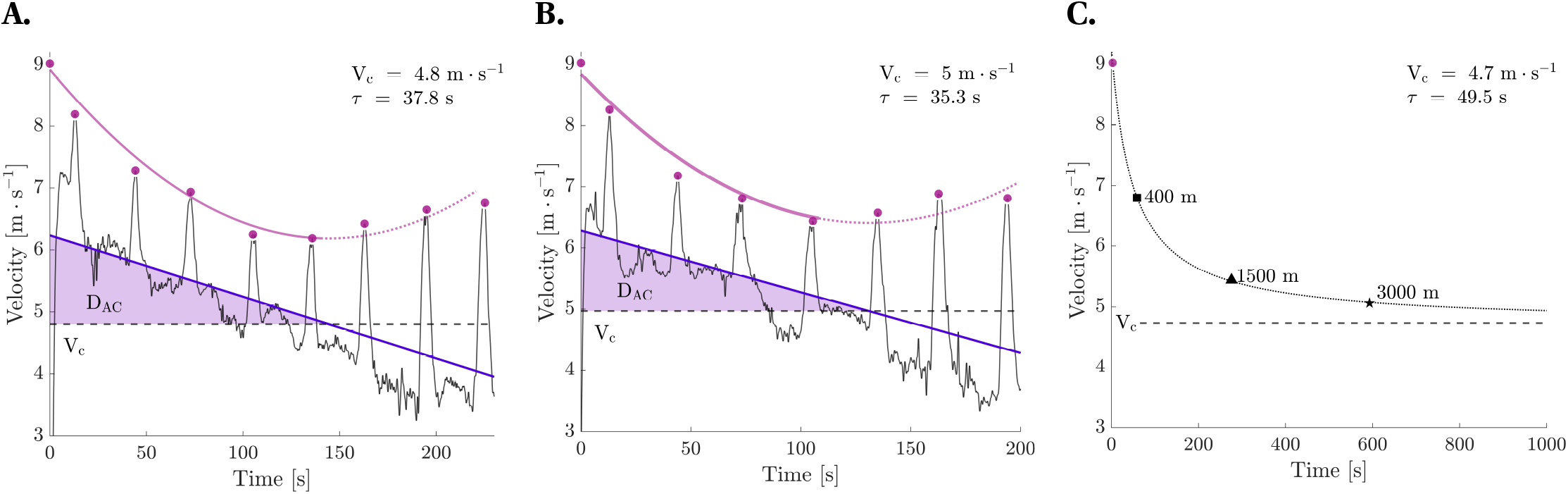
Typical Result: A: RACLET (A), B: RACLET (B), and C: time-trials results for a typical participant (cones setup).

### Feasibility

The mean target velocity was set as *V* ^⋆^ = 5.3 *±* 0.8 m · s^*−*1^ (65.5 ± 7.4 %*V*_*i*_). The median [interquartile range] of the adjusted r2 for Eq. 2 on velocity target of the RACLET test and retest were 0.95 [0.92;0.98] and 0.95 [0.93;0.97] for bike setup and 0.95 [0.89;0.95] and 0.95 [0.90;0.97] for cones setup. The median [inter-quartile range] of r2 were 0.95 [0.93;0.97] and 0.95 [0.93;0.97] for Eq. 8 adjusted on *V*_max_(*t*) of RACLET (A) and RACLET (B) respectively and 0.99 [0.99;0.99] for Eq. 11 adjusted on TT *V*_max_ data. Mean *V*_*i*_ was 8.1 ± 0.7 m · s^*−*1^. The mean perceived exertion of the participants at the most difficult time during the test, and immediately after, was 17.7 *±* 1.7 and 13.1 *±* 2.6 for RACLET (A); 17.1 *±* 1.0, and 13.0 *±* 2.1 for RACLET (B). The mean perceived exertion at the most difficult time and immediately after the test was 16.9 ± 1.9 and 11.6 ± 2.6, respectively, for the bike setup and 18.6 ± 0.9 and 14.7 ± 1.3, respectively, for the cone setup (RACLET (A)). Equivalent values were found for RACLET (B) (16.7 ± 0.9 and 12.2 ± 2.2 for bike setup, and 17.6 ± 0.9 and 14.0 ± 1.5 for cones setup). Significantly higher perceived exertions were observed for cones compared to bike setup (at the most difficult time during the test: p = 0.030, Cohen’s d = 1.09 and p = 0.043, Cohen’s d = 1.00 for RACLET (A) and (B), respectively; at the end of the test: p = 0.006, Cohen’s d = 1.45, and p = 0.049, Cohen’s d = 0.98 for RACLET (A) and (B), respectively).

### Reliability

The differences between RACLET (A) and RACLET (B) parameters are presented in Table. 1. There was no significant difference in the reliabilities of *V*_*c*_, *τ*, and 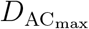 between the bike and cones setups (all p>0.05).

**Table 1.**
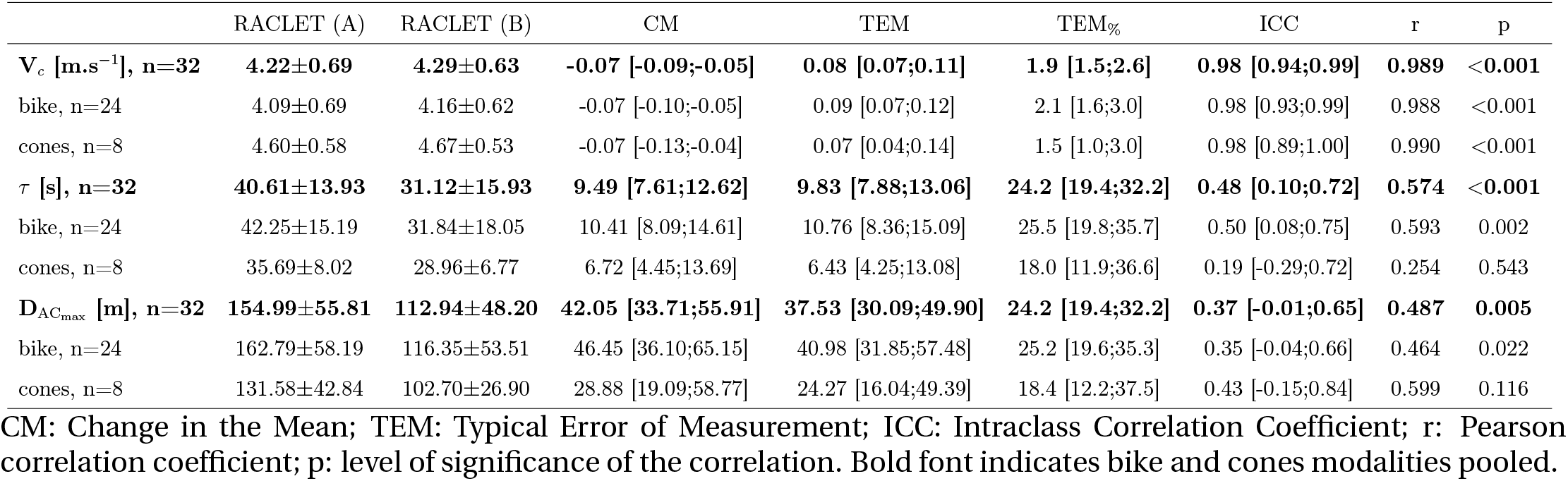
Reliability analysis of the RACLET.

### Validity

There was no significant difference in the validity of *V*_*c*_, *τ*, and 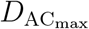 between the bike and cones setups (all p>0.05). The differences between RACLET (A) and TT parameters are presented in Table. 2. Actual and predicted time performances in the 3 TT were not significantly different (p = 0.45, Cohen’s d = 0.11) and were highly correlated (Fig. 4, Table. 3).

**Figure 4.**
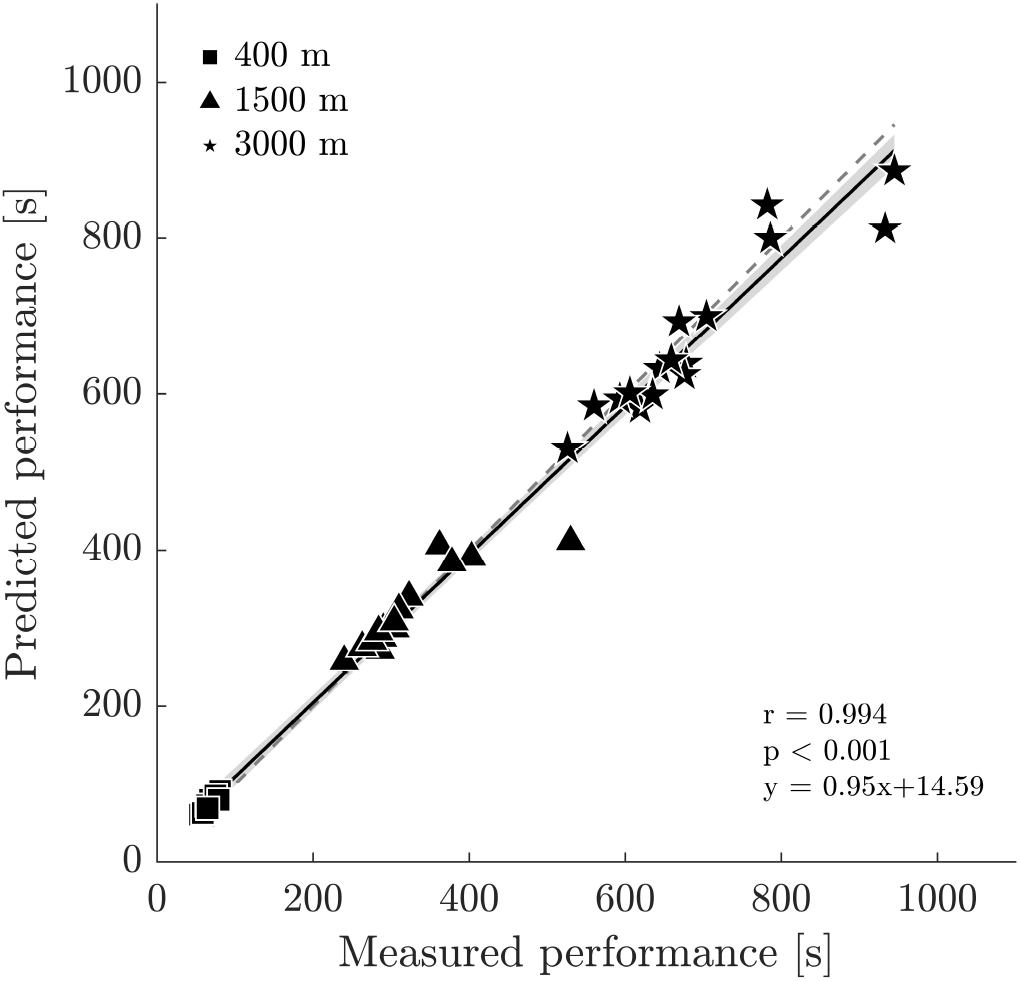
Comparison between predicted and measured time performances on the three time-trials.

## Discussion

The purpose of this study was to test the feasibility of the Ramp Above Critical Level Endurance Test (RACLET), intra-session reliability, concurrent agreement with the time-trials method, and prediction capacity of race time for multiple distances. The main results were (i) an excellent fit of the model to RACLET experimental data (all *r*^2^ *>* 0.89); (ii) an excellent intra-session reliability for *V*_*c*_ (TEM_%_ = 1.9%, ICC = 0.98) but moderate for *τ* (TEM_%_ = 24.2%, ICC = 0.48) and 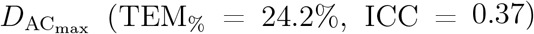; (iii) a good concurrent validity for *V*_*c*_ (SE_%_ ± RE_%_ = 6.2 ± 6.5%) but a significant systematic bias for *τ* (SE_%_ ± RE_%_ = −40.9 ± 29.6%) and 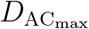 (SE_%_ ± RE_%_ = −43.2 ± 28.9%); (iv) an accurate time-trials performances prediction from parameters determined with the RACLET (SE_%_*±* RE_%_ = 2.60 *±* 7.49%).

First, all participants successfully completed the RACLET, presenting a fatiguing phase that did not lead to exhaustion before starting to recover (see Fig. 3). During the hardest part of the test, a very hard but not maximal effort was perceived on average (17.7 ± 1.7 on the 6-20 Borg scale), in contrast to the time-trial effort which, by definition, leads to maximum effort (19-20; Nicolò et al., 2019). The (active) recovery phase also enabled participants to end the test at a moderate RPE of 13.1 ± 2.6, suggesting the possibility of additional effort (e.g. training) after the test. Furthermore, the experimental observations of maximal velocity assessed from sprints during the RACLET were consistent with the proposed fatigability model; *V*_*max*_(*t*) well followed the expected 2nd order polynomial function (Eq. 10, Fig. 3; adjusted r2 ≥ 0.98). These results are in line with the previous evidences supporting the model’s capacity to describe the fatigability kinetics in the severe domain (Bowen et al., 2024; Vonderscher et al., 2024). Taken together, these results demonstrated that RACLET is highly feasible.

The parameters obtained were in the range of the normative values estimated from the all-out test for similar populations (running athletes in Kramer et al., 2020: *V*_*c*_ ≈ 4 m*·*s*−*1 and 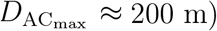. A comparison of *V*_*c*_ between RACLET (A) and (B) showed excellent reliability (ICC = 0.98, TEM_%_ = 1.9%, r = 0.989; Table. 1) within the highest values of ICC range previously reported with various methods (0.919-0.983; Busso et al., 2024; Follador et al., 2020; Nimmerichter et al., 2015; Sousa et al., 2022). Poorer reliability was shown for *τ* (ICC = 0.48, TEM_%_ = 24.2) and for 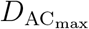 (ICC = 0.37, TEM_%_ = 24.2). Nevertheless, 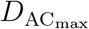 (≡D^*′*^) is known to be associated with a higher random noise than *V*_*c*_ for both time-trials and all-out methods (ICC range 0.325–0.828; TEM_%_ range 12.5 - 26.8 %; Busso et al., 2024; Galbraith et al., 2011; Nimmerichter et al., 2015; Sousa et al., 2022). It should be noted that the 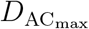 estimated during RACLET (B), conducted 20 min after RACLET (A), was significantly lower, with a moderate effect size (155.0 ± 55.8 vs. 112.9 ± 48.2, p < 0.001; Cohen’s d = 0.75). It is likely that the estimation of 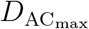 was impacted by RACLET (A) and, unfortunately, incomplete recovery. This was observed to an even greater extent during pilot experiments with a recovery time of 5 min between RACLET, but with no impact on the reliability of *V*_*c*_. These observations indicate that a longer recovery period could contribute to reducing the systematic error and improving the reliability of 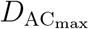, but also that *V*_*c*_ seems still reliably estimated after a fatiguing exercise (e.g. a training session).

RACLET revealed a good concurrent agreement with the time-trials method for critical velocity (r = 0.930; RE_%_ = 6.5), despite a slightly but significantly higher critical velocity for RACLET (SE_%_= 6.2; p < 0.001; Cohen’s d = 0.35). These results are in line with a good agreement, but a general trend toward higher critical velocity estimates for all-out when compared to multiple time-trials or time-to-exhaustion methods (r range 0.695 - 0.93; systematic bias range +2.4 to +10.0%; Aguiar et al., 2018; Broxterman et al., 2013; Hunter et al., 2023). However, larger differences were found for *τ* and 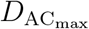 with ≈ 40% lower values for the RACLET compared to the time-trials method (table 2, all p < 0.001; Cohen’s d > 1.5). These large differences in D^*′*^ have already been reported when comparing all-out and time-trials methods (difference range 26-31%; Aguiar et al., 2018; Broxterman et al., 2013; Hunter et al., 2023). Considering *τ*, a direct comparison has never been proposed because it is generally not reported when using the 3-parameter critical speed model. Nevertheless, it is possible to recalculate *τ* from the maximum and critical velocities, as well as D^*′*^: 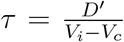. This gives *τ* values of 73.2 to 154.7 s for studies using the time-trials method (Billat et al., 2000; Bosquet et al., 2006; Patoz et al., 2021) whereas the values reported for all-out have a range of 35.2–44.0 s (Kramer et al., 2018, 2020; Kramer et al., 2019; Liu et al., 2021). Interestingly, it has been suggested that all-out does not allow full 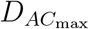 consumption because of pacing strategies, which leads to an overestimation of the critical speed at the end of the test. However, time-trials methods rely conceptually on an outdated definition of *V*_*c*_, which is an intensity that can be maintained “for a very long time without fatigue” (Burnley, 2022). In addition, time-trials are not methodologically designed to estimate *τ* correctly. In contrast, both the RACLET and all-out methods aim to determine *V*_*c*_ as the boundary between the heavy (with a stable physiological state) and severe (associated with drastic fatigability) intensity domains. This could explain why the differences Vonderscher et al. RACLET – Velocity between RACLET and time-trials parameters presented very similar tendencies to those reported between all-out and time-trials methods, raising the question of RACLET concurrent agreement with the all-out method, unexplored in the present study.

**Table 2.** Results of validity between time-trials and RACLET (A).

|  | TT | RACLET (A) | SE | RE | SE<sub>%</sub> | RE<sub>%</sub> | r | p |
| --- | --- | --- | --- | --- | --- | --- | --- | --- |
| **V<sub>c</sub> [m.s<sup>-1</sup>], n=16** | **4.08±0.72** | **4.33±0.64** | **0.25 [0.19;0.39]** | **0.26 [0.20;0.41]** | **6.2 [4.6;9.6]** | **6.5 [4.8;10.0]** | **0.930** | **<0.001** |
| bike, n=8 | 3.75±0.75 | 4.07±0.61 | 0.32 [0.21;0.64] | 0.28 [0.18;0.57] | 8.4 [5.6;17.2] | 7.4 [4.9;15.1] | 0.939 | <0.001 |
| cones, n=8 | 4.41±0.53 | 4.60±0.58 | 0.19 [0.13;0.39] | 0.25 [0.17;0.51] | 4.3 [2.8;8.7] | 5.7 [3.8;11.6] | 0.902 | 0.002 |
| **τ [s], n=16** | **70.97±18.89** | **41.98±14.49** | **-29.00 [-44.88;-21.42]** | **21.01 [15.52;32.52]** | **-40.9 [-63.2;-30.2]** | **29.6 [21.9;45.8]** | **0.229** | **0.393** |
| bike, n=8 | 78.10±21.29 | 48.27±17.18 | -29.83 [-60.70;-19.72] | 26.42 [17.47;53.77] | -38.2 [-77.7;-25.3] | 33.8 [22.4;68.9] | 0.069 | 0.871 |
| cones, n=8 | 63.85±14.00 | 35.69±8.02 | -28.17 [-57.33;-18.62] | 15.69 [10.38;31.94] | -44.1 [-89.8;-29.2] | 24.6 [16.2;50.0] | 0.062 | 0.883 |
| **D<sub>ACmax</sub> [m], n=16** | **278.03±69.56** | **157.88±59.27** | **-120.15 [-185.96;-88.76]** | **80.30 [59.32;124.28]** | **-43.2 [-66.9;-31.9]** | **28.9 [21.3;44.7]** | **0.231** | **0.389** |
| bike, n=8 | 312.70±55.20 | 184.19±64.12 | -128.52 [-261.57;-84.97] | 95.72 [63.29;194.82] | -41.1 [-83.6;-27.2] | 30.6 [20.2;62.3] | -0.283 | 0.497 |
| cones, n=8 | 243.37±67.65 | 131.58±42.84 | -111.79 [-227.52;-73.91] | 67.04 [44.32;136.44] | -45.9 [-93.5;-30.4] | 27.5 [18.2;56.1] | 0.331 | 0.424 |

SE: Systematic error; RE: Random error; r: Pearson correlation coefficient; p: level of significance of the correlation. Bold font indicates bike and cones setups pooled.

More crucially, the validity of the model is also supported by its prediction performance, as tested here through the prediction of duration over 400, 1500, and 3000 m time-trials from the RACLET (A) parameters. Despite the quite low concurrent agreement for 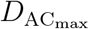 between the RACLET and time-trials methods, excellent prediction accuracy was achieved when forecasting all time-trials performances (−3.5 ± 32.1 s; 2.60 ± 7.49%; r = 0.994, table 3). Only short efforts (of ≈ 60 s) were predicted with mild systematic errors (5.86 ± 3.63 s; 8.85 ± 5.08%; r = 0.945). These results are as good as prediction reported in the literature to estimate time-trials ranging from 800 to 5000 m from the all-out and multiple time-to-exhaustion gold standard methods (typical error range: 2.0 - 15.1 %; r range 0.52-0.98; Bosquet et al., 2006; Busso et al., 2024; Kramer et al., 2018; Laursen et al., 2007; Nimmerichter et al., 2015; Pepper et al., 1992). Predictions based on time-trials and raw in situ data have also shown similarly good accuracy (r = 0.92, in Kranenburg and Smith, 1996; SE_%_ = 7.67%, in Smyth and Muniz-Pumares, 2020).

**Table 3.**
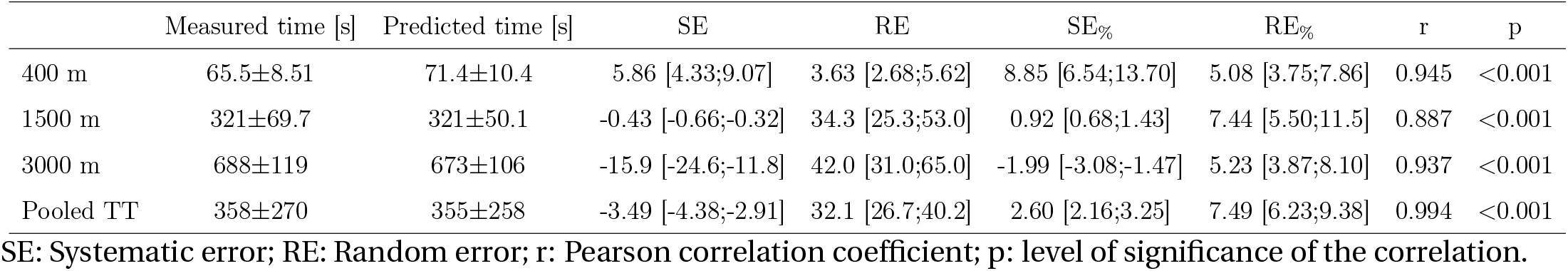
Time trials prediction errors from RACLET (A) parameters (Eq. 12).

|  | Measured time [s] | Predicted time [s] | SE | RE | SE<sub>%</sub> | RE<sub>%</sub> | r | p |
| --- | --- | --- | --- | --- | --- | --- | --- | --- |
| 400 m | 65.5±8.51 | 71.4±10.4 | 5.86 [4.33;9.07] | 3.63 [2.68;5.62] | 8.85 [6.54;13.70] | 5.08 [3.75;7.86] | 0.945 | <0.001 |
| 1500 m | 321±69.7 | 321±50.1 | -0.43 [-0.66;-0.32] | 34.3 [25.3;53.0] | 0.92 [0.68;1.43] | 7.44 [5.50;11.5] | 0.887 | <0.001 |
| 3000 m | 688±119 | 673±106 | -15.9 [-24.6;-11.8] | 42.0 [31.0;65.0] | -1.99 [-3.08;-1.47] | 5.23 [3.87;8.10] | 0.937 | <0.001 |
| Pooled TT | 358±270 | 355±258 | -3.49 [-4.38;-2.91] | 32.1 [26.7;40.2] | 2.60 [2.16;3.25] | 7.49 [6.23;9.38] | 0.994 | <0.001 |

SE: Systematic error; RE: Random error; r: Pearson correlation coefficient; p: level of significance of the correlation.

Overall, the RACLET has the major advantage of being easy to implement. Indeed, the test lasts only 5 min. Two configurations were proposed to follow the velocity target. The first consisted of following a pacing bike, while the second used cones and whistle signals to pace athletes. None of the reliability or validity results were markedly affected by the pacing modality. However, it is interesting to note that the rate of perceived exertion was systematically significantly higher when using cones instead of pacing bike (e.g. RPE = 12.2 vs. 14.0 immediately at the end of RACLET (B) paced with a bike or cones, respectively). This increase in perceived exertion could be due to an increase in cognitive demand when athletes had to self adjust their pace.

However, it still seems easier to implement the cone protocol, particularly when several athletes have to take the test simultaneously or when bikes are not allowed on tracks (or other artificial turf sports pitches).

Further evolutions have already been planned to make the RACLET as easy as possible for both athletes and experimenters. When using cones, the auditive signals could be automated, and the cones were placed 10 m apart to increase feedback resolution. A pacing runner–not performing sprints–could also accompany participants (when feasible) to help with target tracking. Familiarisation is also of paramount importance to help athletes understand the test and to limit their cognitive demands. We suggest performing two familiarisations with at least one execution of the entire test performed at the target velocity sets for the “real” RACLET. Other settings such as the number of sprints could also be modified. However, it should be kept in mind that a good trade-off between a sufficient amount of *V*_max_ evaluations to enable the model adjustment, but not too many sprints to limit fatigability, is preferable. The level of fatigability could also be more controlled by slowing down after sprint phases so that the average velocity during sprint phases would correspond to the target ramp velocity.

The method chosen to set the ramp parameters may evolve in the future. Indeed, it can appear strange that parameters (i.e. *V*_*i*_, *V*_*c*_, and *τ*) should *a priori* be known to choose the ramp settings. However, only rough approximations are needed to ensure a feasible test. The parameters obtained *a priori* are only used to distinguish endurant (runners) from the more explosive (e.g. team sport athlete) profiles. From these rough evaluations of the parameters, V^⋆^ was determined to induce a depletion of 50% of the 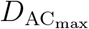. However, the actual *D*_AC_ consumed during the RACLET ranged between 13.7 and 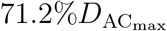. This difference clearly indicates that the rough *V*_*i*_, *V*_*c*_, and *τ* estimations are close to, but not accurately representative of, the participants’ capacities. The same settings can also be used as a basis to re-test athletes with possible adjustments to anticipate improvements (for example, after training). If the athlete capacities have improved, it is likely that t_*c*_ will appear earlier in the test–and so that the test will lead to less fatigue–resulting in a greater *V*_*c*_ for a similar ramp setting. *Nota bene*, for a given athlete, at a given time, various ramp settings should theoretically result in similar output parameters (*V*_*i*_, *V*_*c*_, and *τ*), although this remain to be tested experimentally.

Due to its simplicity, the RACLET offers new perspectives that deserve to be mentioned. Indeed, the easiness-to-setup and shortness of the test (5 min) enable routine testing of athletes’ capacities, even within a training session. In addition, future RACLET can be adapted to account for the specific demands of team sports. Target tracking could be performed in shuttle running between two cones (40-m apart), therefore incorporating changes of direction and mimicking the multidirectional movement patterns typical in team sports. From a research perspective, the RACLET paves the way to original designs aimed at evaluating the effects of any factor (e.g. hill gradient) on the critical velocity, therefore requiring several critical velocity tests that are not easy to implement with historical methods, e.g. in a trail running context.

Finally, it is crucial to remember that all the results presented here are directly related to the quality of the experimental protocol (in both settings and measurements). The authors kindly encourage any person intending to use this test to focus on all the methodological aspects. An improperly configured test, whether too easy or too difficult, bad target tracking, submaximal efforts during sprint phases, or velocity measurement of poor quality are likely to affect reliability and validity. To assist anyone interested in implementing the RACLET, all the procedures for designing and processing the RACLET, along with extensive explanations were made available in Excel and Matlab formats on the FoVE research group’s Gitlab: https://gricad-gitlab.univ-grenoble-alpes.fr/fove/methods/raclet.

## Conclusion

This study demonstrated that a simple five-minute decreasing ramp test (the Ramp Above Critical level Endurance Test, RACLET) based on a mathematical model of fatigability is reliable and valid for determining the critical velocity during running. Its predictive capacity is in line with that of historical tests, with the major advantage of not leading to exhaustion. The test was very well tolerated by all participants, confirming its potential for regular use in the field (e.g. to monitor training) or to address scientific questions covering multiple conditions (e.g. the effect of hill gradients on critical velocity).

## Abbreviations

*D*_AC_: Distance covered Above Critical velocity
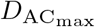: Maximal Distance covered Above *V*_*c*_
RACLET: Ramp Above Critical Level Endurance Test
*S*: Slope of the RACLET ramp
*τ*: Typical time of fatigability
*t*_*c*_: Time at which Critical velocity is reached
*t*_lim_: Time of exhaustion (variable)
*t*_*r*_: Total duration of RACLET Ramp
*t*_*T*_: Time-trial duration
*TT*: Time-trial (method)
*V*_*i*_: Initial maximal theoretical velocity
*V*_*c*_: Critical theoretical velocity
*V*_end_: Velocity at the end of the RACLET
*V*_max_: Maximal theoretical velocity
*V* _⋆_: Velocity at the start of the RACLET

## Acknowledgments

The authors would like to thank Christian Osgnach and the GPexe team for allowing the use of their equipment in this study. Support was provided by the French National Agency [ANR-22-CE14-0073] to B. Morel, and by the Conseil Savoie Mont Blanc (CSMB) to M. Vonderscher. The authors have no conflicts of interest to disclose relevant to the content of this article.

